# Optical modulation of excitation-contraction coupling in human induced pluripotent stem cell-derived cardiomyocytes

**DOI:** 10.1101/2022.09.22.508828

**Authors:** Vito Vurro, Beatrice Federici, Carlotta Ronchi, Chiara Florindi, Valentina Sesti, Silvia Crasto, Claudia Maniezzi, Camilla Galli, Maria Rosa Antognazza, Chiara Bertarelli, Elisa Di Pasquale, Guglielmo Lanzani, Francesco Lodola

**Author notes:** Corresponding authors: Francesco Lodola; Guglielmo Lanzani. Eindhoven University of Technology, 5612 AZ Eindhoven, Netherlands (BF); Amarin Italy S.r.l., Corso Vercelli 40, 20145, Milan, Italy (SC).

## Abstract

Non-genetic photostimulation is a novel and rapidly growing multidisciplinary field of research that aims to induce light sensitivity in living systems by exploiting exogeneous phototransducers.

Here we propose a recently synthetized intramembrane photoswitch, based on an azobenzene derivative (Ziapin2), for optical pacing of human induced pluripotent stem cell-derived cardiomyocytes (hiPSC-CMs).

The light-mediated stimulation process has been studied by applying several characterization techniques to detect the effect on the cell properties. In particular, we recorded changes in membrane capacitance, in membrane potential (V_m_), and modulation of intracellular Ca^2+^ dynamics. Finally, cell contractility was analyzed using a custom MATLAB algorithm.

Photostimulation of intramembrane Ziapin2 causes a transient V_m_ hyperpolarization followed by a delayed depolarization and action potential firing. The observed initial electrical modulation nicely correlates with changes in Ca^2+^ dynamics and contraction rate.

This work represents the proof of principle that Ziapin2 can modulate electrical activity and contractility in hiPSC-CMs, opening up a future development in cardiac physiology.

## 1. Introduction

In the cardiovascular field, optical stimulation is emerging as an alternative to traditional approaches for many research and therapeutic applications thanks to a series of key-enabling features: the lower energy consumption and release, minimal invasiveness net of an extraordinary spatial and temporal resolution (Antognazza et al., 2015; Di Maria et al., 2018; Hopkins et al., 2019).

Optogenetics (Deisseroth, 2011; Fenno et al., 2011; Rein and Deussing, 2012; Deisseroth, 2015; Duebel et al., 2015; Dwenger et al., 2019), that has become widespread in neuroscience, could in principle be relevant in the cardiac field too (Knollmann, 2010; Bruegmann et al., 2010; Ambrosi et al., 2014; Boyle et al., 2018; Asano et al., 2018; Ferenczi et al., 2019; Sasse et al., 2019; Krueger et al., 2019; Huang et al., 2020; Floria et al., 2020; Li et al., 2021; Crocini et al., 2016; Biasci et al., 2022). However, the approach still has a limited clinical applicability mainly because cell optical sensitivity is obtained by transduction with gene constructs carried by viral vectors.

A possible alternative strategy to overcome these constraints relies on the use of light-sensitive transducers, based on both inorganic and organic semiconductors (Starovoytov et al., 2005; Bareket-Keren and Hanein, 2014; Savchenko et al., 2018; Rand et al., 2018; Feiner and Dvir, 2019; Lodola et al., 2019a, 2019b; Parameswaran et al., 2019; Bruno et al., 2021; Negri et al., 2022). Those have been used recently as photoactive interfaces for cardiomyocyte (CM) optical stimulation. The generation of action potentials (APs) has been demonstrated by using planar graphene-based biointerfaces, possibly exploiting the photogeneration of charge carriers, and with polymer-silicon nanowires utilizing their photoelectrochemical properties (Savchenko et al., 2018; Parameswaran et al., 2019). Similar results were obtained also increasing the local temperature with illumination of absorbers in contact with CMs, using gold nanoparticles or nanorod electrodes and metasurface planar organic interfaces (Gentemann et al., 2017; Bruno et al., 2021; Fang et al., 2022). Organic semiconductors in particular allow to establish almost seamless abiotic/biotic interfaces, as they are soft materials that support both ionic and electronic transport, alike many biological molecules (Rivnay et al., 2014). In this context, the triggering mechanism leading to cell activity upon light absorption can be capacitive, faradaic, or thermal (Martino et al., 2015; Ðerek et al., 2020; Manfredi et al., 2021).

An alternative approach, still based on organic molecules, exploits photochromic compounds that can be covalently bound (Gorostiza and Isacoff, 2008; Izquierdo-Serra et al., 2016; Leippe et al., 2017) to an ion channel or not-covalently (Zhang et al., 2014; Fang et al., 2020) to the plasma membrane. This methodology is gaining increasing interest due to the stimulation efficiency, its versatility and the possibility of using two photon absorption that pushes the stimulation wavelengths to the near infrared spectral region, improving tissue penetration (Magome et al., 2011; Riefolo et al., 2019).

In this work, we propose a new tool for optical pacing of human-induced pluripotent stem cell-derived CMs (hiPSC-CMs) exploiting Ziapin2, a recently synthetized intramembrane photochromic transducer that we successfully tested in bacteria, non-excitable cells and neurons (DiFrancesco et al., 2020; Paternò et al., 2020a, 2020b; Vurro et al., 2021; Magni et al., 2021).

Ziapin2 has affinity for the hiPSC-CMs sarcolemma and once partitioned, it undergoes trans-dimerization, which in turn leads to increased capacitance as the result of reduction in membrane thickness. Upon millisecond pulses of visible light, *trans→cis* isomerization causes a fast drop of capacitance due to membrane relaxation, resulting in a transient hyperpolarization followed by a delayed depolarization that triggers AP generation. The increase in AP frequency nicely correlates with changes in Ca^2+^ dynamics and contraction rate, thus proving that Ziapin2 modulates excitation-contraction (E-C) coupling at a whole extent.

## 2. Results and Discussion

### 2.1. Ziapin2 photoisomerization modulates hiPSC-CMs membrane potential

The molecule was tested at two different concentrations (5 μM and 25 μM). In both cases once added to hiPSC-CMs cultures, Ziapin2 successfully partitioned into the sarcolemma causing a significant increase in capacitance (**Figure S1a**). The effect was concentration dependent, as we observed a +23% rise (14.2 ± 1.2 pF vs 17.5 ± 0.9 pF; p = 0.05089) for 5 μM and +56% (14.2 ± 1.2 pF vs 22.2 ± 1.3 pF; p = 0.00063) for 25 μM, while no alterations were detected in hiPSC-CMs incubated with the vehicle (DMSO, data not shown). At equilibrium, this phenomenon changes the membrane electrical time constant (RC) but not the resting potential. According to previous experimental evidences (DiFrancesco et al., 2020; Paternò et al., 2020b; Vurro et al., 2021), the modulation has been ascribed to the thinning of the bilayer caused by the dimerization of Ziapin2 molecules inside the membrane.

Upon photostimulation the azobenzene isomerization removes the geometric constraint of the dimer leading to membrane relaxation and a partial return towards steady-state capacitance values with a reduction of −14% (17.5 ± 0.9 pF vs 15.0 ± 0.7 pF; p = 0.08) and −12% (22.2 ± 1.3 pF vs 19.5 ± 1.1 pF; p = 0.14) for 5 and 25 μM Ziapin2, respectively (**Figure S1b**).

Whole-cell patch clamp experiments in current clamp mode (I=0) revealed that the photoinduced drop in capacitance correlates with a modulation of the membrane potential. In particular, the Ziapin2-mediated photostimulation (with 20 and 200 ms single light pulses) results in a transient hyperpolarization that is occurring within few ms after the light onset, consistent with the linear inverse relation between capacitance and voltage. The recovery of the potential by capacitive currents is followed by a depolarization which is often large enough to generate an AP (see representative traces in **Figure 1a**).

**Figure 1.**
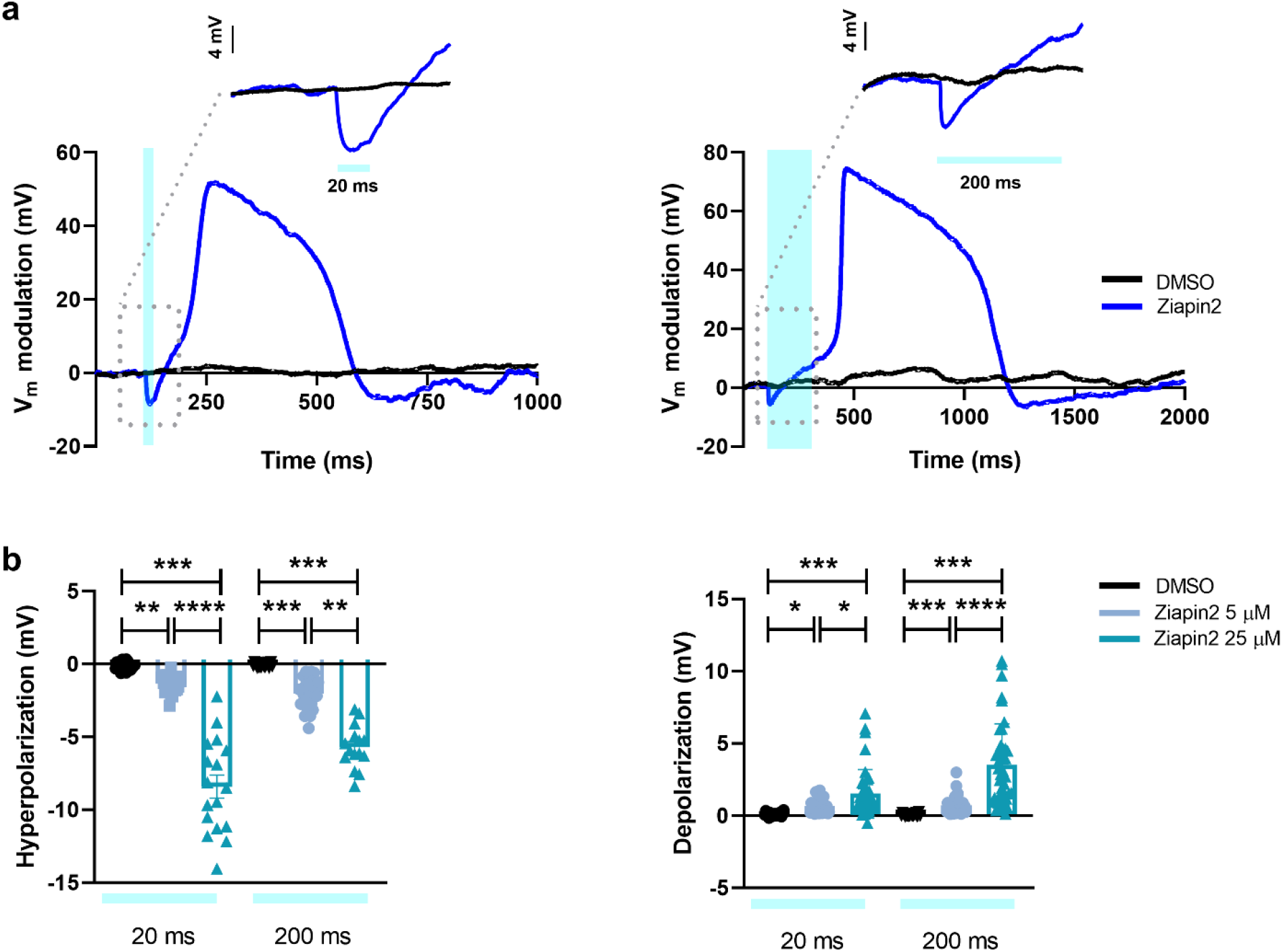
Ziapin2 mediates a light-evoked membrane voltage modulation in hiPSC-CMs. **a)** Representative whole-cell current-clamp traces recorded in cells loaded with either vehicle (DMSO, in black) or Ziapin2 (blue) and stimulated with 20 ms- (left) or 200 ms-long (right) single light pulses. Photoexcitation is represented by the cyan shaded area. Light power density, 80 mW/mm^2^. The membrane potential modulation can be appreciated at a faster time scale in the inset of each panel. **b)** Scatter plots of the peak hyperpolarization (left) and depolarization (right) changes in hiPSC-CMs exposed to Ziapin2 (5 μM in sky blue, 25 μM in teal) or DMSO for the above-mentioned light-stimulation protocols. n >18 for 5 μMand 25 μM Ziapin2 loaded cells and n = 20 for DMSO-treated hiPSC-CMs. V_m_ values have been reported as relative variation to better appreciate the light-induced effect; however, no significant changes were detected in the resting membrane potential values (Ziapin2 5μM V_m_ = −42.4 ± 3.9 mV; Ziapin2 25μM V_m_ = −42.8 ± 2.6 mV; DMSO V_m_ = −37 ± 4.4 mV). The experiments were carried out at room temperature (24°C). Data were collected from three independent differentiations and are represented as mean ± sem. * p < 0.05, ** p < 0.01, *** p < 0.001 and ****p < 0.0001.

Both hyperpolarization and depolarization peaks display much higher amplitude at 25 μM concentration than 5 μM (**Figure 1b**). For instance, with 20 ms light stimulation hyperpolarization is six times larger at 25 μM concentration, even if the relative change in capacitance is comparable (−14% vs −12% for 5 and 25μM Ziapin2, respectively, see **Figure S1b**). This is surprising, since according to the simple plane capacitor model, if there are no other changes in the circuit, ΔV/V=ΔC/C. To explain this discrepancy, we conjecture that equal amplitude, but faster changes in capacitance, could give rise to larger changes in membrane potential. This is confirmed by a numerical simulation based on the equivalent circuit model for the membrane (**Figure S2**). To account for the different kinetics, we propose that at higher concentration, Ziapin2 dwells in a more heterogeneous membrane environment, characterized by different viscosity, order and phospholipid polarity, where a faster isomerization might occur.

In agreement with capacitance data, no light-dependent effects were noticed in vehicle-treated hiPSC-CMs (**Figure 1**).

### 2.2. Ziapin2 photoisomerization induces AP firing in hiPSC-CMs

To assess the impact of the molecule on hiPSC-CMs electrical activity, we evaluated the ability of light stimulation to generate APs in hiPSC-CMs loaded with Ziapin2. Remarkably, the most striking effect was observed in CMs exposed to the highest photochromic concentration (72 ± 4% of responding cells for Ziapin2 25 μM vs 37 ± 3% for Ziapin2 5 μM, **Figure 2a**).

**Figure 2.**
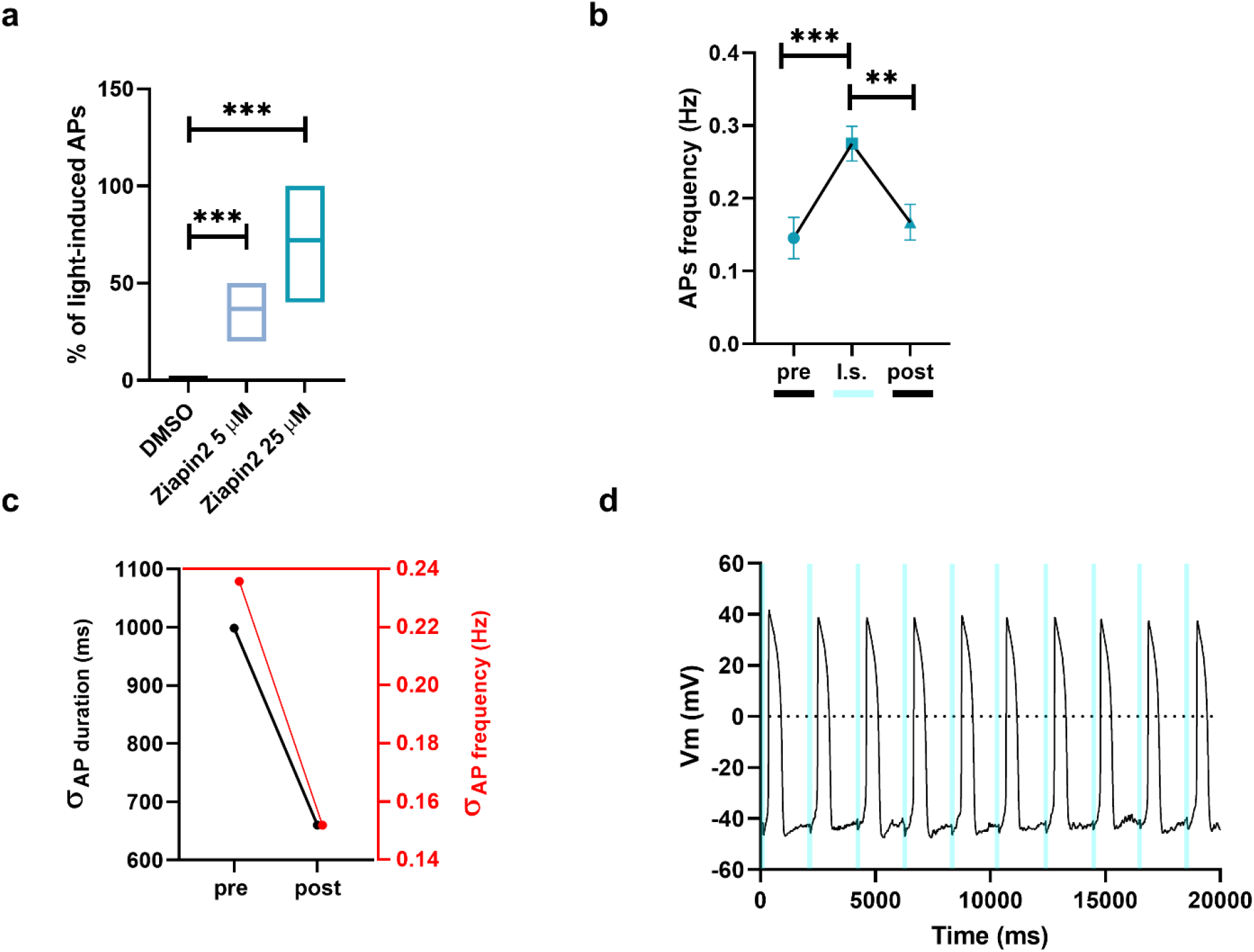
Light-evoked AP modulation by Ziapin2 in hiPSC-CMs. **a)** Box plots (line at mean) of the reproducibility of the light-induced AP generation in cells loaded with either vehicle (DMSO, in black) or Ziapin2 (5 μM, in sky blue; 25 μM, in teal). The percentage represents the n° of AP directly generated by each light pulse. **b)** AP frequency before (pre), during (l.s.), and after (post) light stimulation. Data are represented as mean ± sem; n > 40 for each condition from three independent differentiations. **c)** Correlation between the standard deviation (σ) of AP duration (left, black) and frequency (right, red) pre- and post-photostimulation. **d)** Representative traces of APs generated in response to a light train stimulation (20 ms cycle length; 0.5 Hz stimulation frequency, represented as cyan shaded areas). Light power density, 80 mW/mm^2^. The experiments were carried out at room temperature (24°C). **p < 0.01 and ***p < 0.001.

As a next step, we compared spontaneous and light-evoked AP features. Since the most significant effects occurred in hiPSC-CMs loaded with Ziapin2 25 μM and subjected to 20 ms light stimuli, here we report only data within these experimental conditions.

In dark conditions no significant differences were measured in maximum diastolic potential (MDP, **Figure S3a**) and maximal upstroke velocity (dV/dtmax, **Figure S3b**), while AP amplitude (APA, **Figure S3c**) was slightly decreased in Ziapin2-exposed CMs (70.3 ± 1.2 mV vs 70.8 ± 2.9 mV; p = 0.04). Ziapin2 photoisomerization did not exert any detectable modification of these AP parameters (**Figure S3b and S3c**), while a light-triggered increase in the number of AP per unit time was observed (0.14 ± 0.02 Hz vs 0.27 ± 0.02 Hz for spontaneous and light-evoked respectively; p < 0.0001, **Figure 2b**). This light-induced frequency increment was associated with a shortening of the AP duration; plotting the standard deviation of these two parameters, it was possible to notice a greater clustering of both the indicators upon illumination (**Figure 2c)**. This suggests that Ziapin2 can potentially play a pivotal role in making homogeneous activity of cardiac cells and tissues by reducing the overall beating variabilities. Interestingly, a punctual AP generation was obtained when a stimulation train of 20 ms light pulses at 0.5 Hz was applied, observing just few failures under rate-controlled conditions (**Figure 2d**). We attributed the missed stimulations to the low temperature (24°C) used in our experiments. It is indeed known that a more physiological temperature would speed up the dynamics of cellular processes. Accordingly, we monitored the electrical activity of hiPSC-CMs at 37°C (**Figure S4**). As expected, in these conditions there was a higher spontaneous electrical activity (~ 0.78 Hz, **Figure S4a, left panel**) that increased upon Ziapin2-mediated photostimulation (~ 0.95 Hz, **Figure S4b, left panel**). Notably, considering a sampling time of 2 seconds, we observed that the APs under control conditions are randomly distributed (**Figure S4a, right panel**), while upon photoexcitation they cluster at the light onset (**Figure S4b, right panel**). This means that the spontaneous APs are generated in deterministic way and that we are imposing a 0.5 Hz stimulation frequency.

From this small subset of data, we also extrapolate the AP duration at 90% of repolarization (APD90, directly correlated to the frequency increase) and APA. In **Figure S4c** are reported data acquired at both 24° and 37°C to show that, as expected, the increase in temperature mainly affects APD90, leaving the amplitude unaltered.

### 2.3. Ziapin2 photoisomerization elicits Ca^2+^ dynamics in hiPSC-CMs

In CMs the AP triggers the E-C coupling, namely the physiological process that converts an electrical stimulus to a mechanical response. It is well established that the contraction of cardiac cells is triggered by a considerable increase in intracellular Ca^2+^ concentration. We thus evaluated Ziapin2 effect on cytoplasmic Ca^2+^ levels performing fluorescence-based measurements with the widely used ratiometric Ca^2+^ indicator Fura-2 AM (**Figure 3a**).

**Figure 3.**
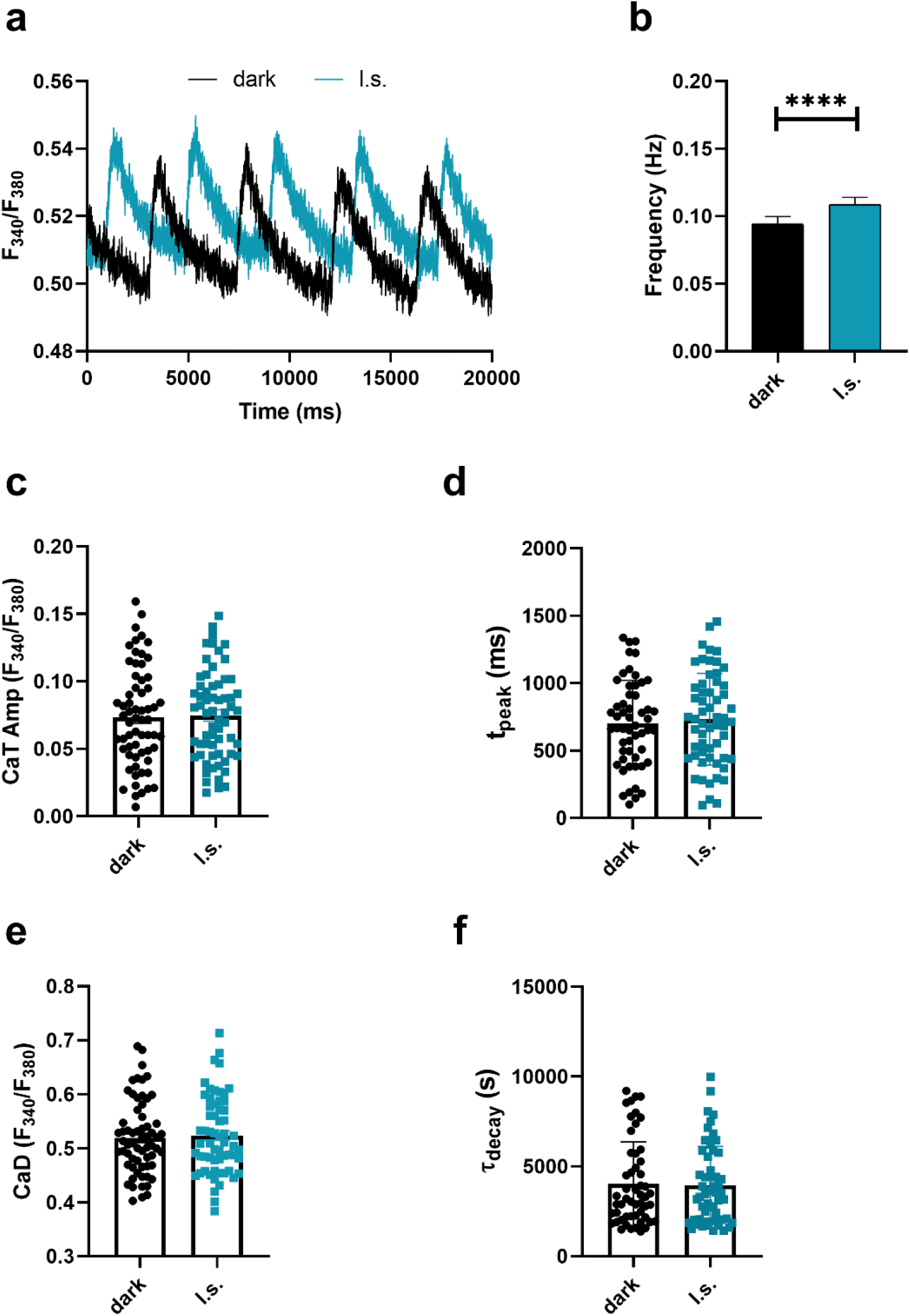
Photo-induced Ca^2+^ transients in Ziapin2-loaded hiPSC-CMs. **a)** Representative Ca^2+^ transients (CaTs) recorded on the same Ziapin2-loaded cell first in dark condition (dark, in black) and subsequently photostimulated with a 200 ms pulse (l.s., in teal) at light power density = 273 μW/mm^2^. **b)** Frequency of CaTs, **c)** CaT amplitude (CaT Amp), **d)** CaT rise-time (t_peak_), **e)** Diastolic Ca^2+^ (CaD), **f**) CaT decay kinetics (τ_decay_). The paired measurements were carried out at room temperature (24°C) on 63 cells. ****p < 0.0001.

To monitor if the molecule photoisomerization had an impact on Ca^2+^ dynamics, we first measured the spontaneous Ca^2+^ activity by keeping Ziapin2-loaded hiPSC-CMs in dark conditions, and subsequently, we subjected the same cells to a single light stimulation pulse using an arc lamp light source coupled with a high-speed OptoScan monochromator that allowed us to accurately select the excitation wavelength.

In accordance with the patch-clamp data we observed a significantly higher frequency of Ca^2+^ transients (CaTs) (+15.4%, p = 0.0001, **Figure 3b**) when hiPSC-CMs were subjected to light stimulation while no significant changes were detected in the amplitude (p = 0.6, **Figure 3c**), in the rise-time (p = 0.93, **Figure 3d**), nor in the decay kinetics (p = 0.67, **Figure 3e**) of the CaTs. Moreover, the diastolic Ca^2+^ levels were also comparable among the two groups (**Figure 3f)**.

Since changes in the shape of the CaT might reflect alterations in cardiac ion channels or cardiac signaling pathways, we can speculate that is unlikely that a primary membrane transporter like Na^+^/Ca^2+^ exchanger (NCX) or the plasma membrane Ca^2+^ ATPase (PMCA) are altered upon Ziapin2 photoisomerization, suggesting that the molecule acts by fully respecting the physiological behavior of the cell.

### 2.4. Ziapin2 photoisomerization enhances hiPSC-CMs contraction rate

As a final step, we investigated the effect of Ziapin2 on hiPSC-CMs contraction behavior by exploiting a custom MATLAB code that allowed us to obtain a quantitative analysis from previously acquired high-speed movies **(Figure 4)**.

**Figure 4.**
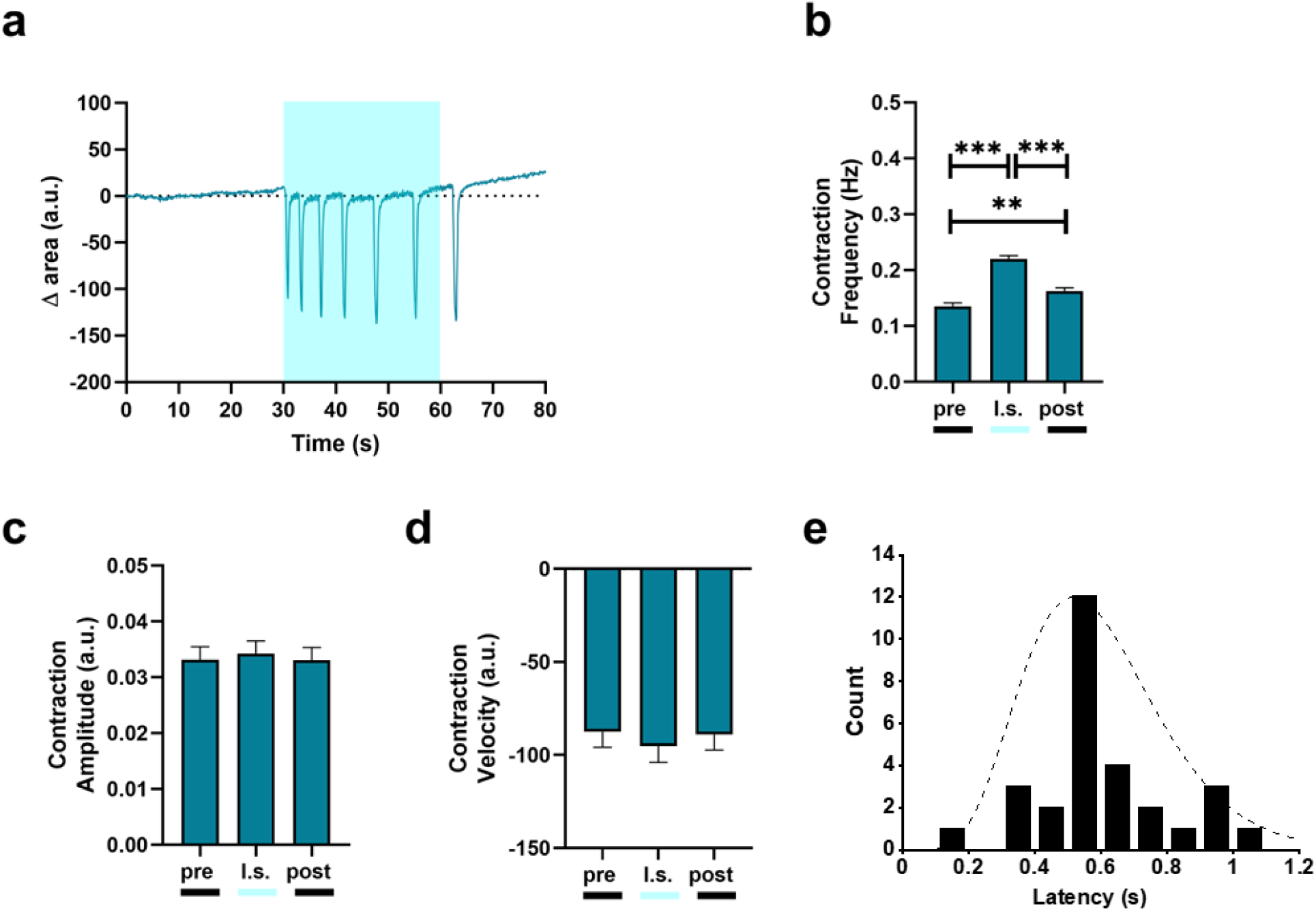
Light-induced contraction rate modulation in Ziapin2 loaded hiPSC-CMs. **a)** Representative trace of the contraction behaviour of Ziapin2-treated hiPSC-CMs. **b)** Contraction frequency, **c)** Contraction amplitude, and **d)** Contraction velocity analyzed before (pre), during (l.s.), and after (post) 1 Hz pulsed light stimulation at light power density = 30 mW/mm^2^ (n = 220 from three independent differentiations, data are shown as mean ± sem). **e)** Time to contraction measured from the light onset in quiescent hiPSC-CMs (n = 29). The cell response time distribution peaks at 500 ms, highlighting a correlation between light stimulation and cell’s functional response. A gamma function has been used as a guide to the eye of this time distribution. The experiments were carried out at room temperature (24°C). **p < 0.01 and ***p < 0.001.

In dark condition, Ziapin2-loaded hiPSC-CMs had a contraction rate of 0.13 ± 0.006 Hz. Under 1 Hz pulsed illumination the rate increased by 59% during the entire acquisition window (**Figure 4b**), while it remained constant in vehicle-treated hiPSC-CMs subjected to the same stimulation protocol (**Figure S5**). The augmented frequency occurs net of variations in contraction amplitude (calculated as area reduction, p = 0.74, **Figure 4c**) and contraction velocity (extracted as derivative of the contraction amplitude, p = 0.57, **Figure 4d**). Interestingly, the effect on the contraction frequency persisted in the 25-30 seconds following light offset, albeit with a 27% decrease (from 0.22 ± 0.006 Hz to 0.16 ± 0.0059 Hz, **Figure 4b**). There are two possible explanations for this; first, the experiments were performed at 24°C, a temperature far from the canonical physiological 37°C that could have slowed down the cell behavior (Lodola et al., 2019b). Secondly, even though the hiPSC-CMs are a consolidated tool and hold an enormous potential in cardiovascular research, their E-C coupling process resembles more the one of fetal/neonatal CMs, thus it is possible that the Ca^2+^ clearance machinery in these cells is substantially slower in comparison to adult CMs (Kane et al., 2015; Lodola et al., 2021).

Remarkably, the lower the spontaneous hiPSC-CMs contraction rate, the higher the light induced frequency increase. To better characterize the involvement of Ziapin2 in the light-induced increase in contraction frequency, we measured the latency, calculated as the interval between the onset of stimulation and the contraction of the CMs, in a small subset of quiescent cells (**Figure 4e**).

The timescale was found to be distributed around 500 ms in a non-stochastic way, compatibly, at least in these experimental conditions, with expected times for the electromechanical coupling, thus confirming the link between optical stimulus and increased contraction.

## 3. Conclusions & Perspectives

The possibility to manipulate the cellular activity with a precise spatial and temporal punctuality is a key enabling technology for biology and medicine. In this work we report the first proof-of-principle demonstration of Ziapin2 as a light-sensitive tool to control cardiac cell excitability and contractility. Our approach is innovative for a series of reasons: i) unlike electrical stimulation, it enables a non-contact and reversible excitation with high spatio-temporal precision; ii) it avoids genetic manipulation and the pitfalls related to the introduction of exogenous genetic material using viral vectors, thus overcoming the major drawback of optogenetics; iii) Ziapin2 primarily targets the cellular passive properties without affecting ionic conductances, an issue shared by both optogenetics and photoswitchable ligands, still allowing a millisecond control of V_m_; iv) the molecule effect is heatless, thus preventing increases in temperature that could be potentially harmful to the cells over the long term.

Ziapin2 could be used as a non-invasive optical tool to control cardiac electrical activity for bio-hybrid robotics studies (in which CMs are often the biological substrate), thus enabling or simplifying design and fabrication of the actuators (Park et al., 2016; Raman et al., 2016; Vurro et al., 2022). Also, the possibility to pattern light in cardiac organoids or microtissues will allow to address fundamental questions of cardiac biology.

## 4. Limitations of the study

Further studies are needed to elucidate the biophysical mechanisms ruling the photostimulation process. In particular, future research will be devoted to the study of the triggering mechanism that leads to AP generation and to exclude the potential involvement of membrane transporters like NCX, Na^+^ or K^+^-ATPase (NKA) as well as membrane receptors (e.g., β-Adrenergic receptors). Moreover, some chemical improvement should be faced to increase the molecular absorption cross section leading to a reducing of the required light power density. Similarly, to further corroborate our findings, we will extend this approach to fully differentiated, adult cardiomyocytes, either human- or mouse-derived. This will be an important step to show that our molecules do work in a more complex and physiological scenario.

If successful, this could open a broader field of applications for light-mediated clinical treatment of cardiac arrhythmias. In this context, new curative strategies that are less invasive, long-lasting, and less power consuming than the existing ones are a *conditio sine qua non* to reduce the burden for the recipients.

## 5. Star Methods

Detailed methods are provided in the online version of this paper and include the following:

- **Key Resources Table**
- **Resource Availability**
  - **Lead Contact**
  - **Materials availability**
  - **Data and code availability**
- **Experimental model and subject details**
- **Method Details**
- **Statistical Analysis**

## 6. Supplementary Information

Supplemental information can be found online.

## 7. Acknowledgements

The authors would like to thank Prof. Antonio Zaza for the interesting and inspiring discussions on this work. This research was supported by: Fondo di Ateneo (2020-ATE-0044) from the University of Milano-Bicocca (F.L.), Nanosparks project (ID 2018–0505) from Fondazione Cariplo (G.L.), EU Horizon 2020 FETOPEN-2018-2020 Programme ‘LION-HEARTED’, grant agreement n. 828984 (M.R.A., F.L., C.R., C.G., E.D.P.). Graphical Abstract created with BioRender.com.

## 8. Author contributions

Vito Vurro: Conceptualization, Data curation, Formal analysis, Investigation, Writing – review & editing; Beatrice Federici: Data curation, Formal analysis, Investigation; Carlotta Ronchi: Data curation, Formal analysis, Investigation; Chiara Florindi: Data curation, Formal analysis, Investigation; Valentina Sesti: Investigation, Methodology; Silvia Crasto: Investigation, Methodology; Claudia Maniezzi: Data curation, Formal analysis; Camilla Galli: Investigation, Methodology; Maria Rosa Antognazza: Investigation, Methodology, Supervision; Chiara Bertarelli: Investigation, Methodology, Supervision; Elisa Di Pasquale: Investigation, Methodology, Supervision; Guglielmo Lanzani: Conceptualization, Funding acquisition, Investigation, Resources, Supervision, Validation, Writing –review & editing; Francesco Lodola: Conceptualization, Funding acquisition, Investigation, Project administration, Supervision,Validation, Writing – original draft.

## Star Methods

### Key Resources Table

3 Key Resources Table
| REAGENT or RESOURCE | SOURCE | IDENTIFIER |
| --- | --- | --- |
| **Biological samples** |  |  |
| hiPSC-CMs | Ncardia | Pluricyte® Cardiomyocyte, PCMI-1033-1 |
|  | Humanitas Clinical and Research Center-IRCCS | See below for the differentiation protocol. |
| **Chemicals** |  |  |
| Fura-2-AM | ThermoFisher Scientific | F1201 |
| hiPSC growth medium | ThermoFisher Scientific | StemFlex Medium (A3349401) |
| Vitronectin (VTN-N) Recombinant Human Protein, Truncated | ThermoFisher Scientific | A14700 |
| hiPSC-CMs medium | Ncardia | Pluricyte® Cardiomyocyte Medium, PM-2100-100mL |
| B27-Supplement w/o insulin | ThermoFisher Scientific | 0050129SA |
| B27 Supplement | ThermoFisher Scientific | 17504044 |
| Growth Factor-Reduced Matrigel (10ml) | BD Biosciences | 354230 |
| RPMI Medium 1640 with L-Glutamine | ThermoFisher Scientific | 11875-119 |
| CHIR-99021 | Selleckchem | S1263-5MG |
| IWR1 | Sigma Aldrich | 10161-5MG |
| Y-27632 (ROCK Inhibitor) | Selleckchem | S1049 |
| Laminin | Sigma Aldrich | L2020 |
| Fibronectin | Sigma Aldrich | F0895-2MG |
| **Software and algorithms** |  |  |
| Microsoft Excel | Microsoft | https://www.microsoft.com/ |
| GraphPad Prism | Prism | https://www.graphpad.com/scientific-software/prism/ |
| MATLAB | MathWorks | https://it.mathworks.com/products/matlab.html |
| Origin 9.0 | OriginLab Corporation | https://www.originlab.com/index.aspx?go=Support&pid=1960 |
| pClamp-10 software<br>Contraction analysis code | Axon Instruments | http://go.moleculardevices.com/l/83942/2015-09-08/77t9w |
|  | This paper | https://github.com/BeatriceFederici/contraction_tracking.git |
| **Other** |  |  |
| Patch Clamp Setups | Molecular Devices | Axopatch 200B, Digidata 1440A<br>MultiClamp 700B, Digidata 1440A |
| Microscopes | Nikon | Eclipse Ti<br>Eclipse TE200 |
| Illumination sources | Lumencor<br>Cairn Research | Spectra X<br>Arc Lamp Light Source coupled with high-speed OptoScan monochromator |
| Cameras | Photometrics<br>ThorLabs | Prime 95B Scientific CMOS (sCMOS) Camera<br>CS135MUN, Kiralux 1.3 MP NIR-Enhanced CMOS Camera |

#### Lead Contact

Further information and requests for resources and reagents should be directed to and will be fulfilled by the lead contacts, Francesco Lodola and/or Guglielmo Lanzani.

#### Materials Availability

This study did not generate any unique new reagents.

#### Data and code availability

- All the data reported in this paper will be shared by the lead contact upon request.
- All original code has been deposited to Github and is publicly available as of the date of publication. The DOI is listed in the key resources table.
- Any additional information required to re-analyze the data reported in this paper is available from the lead contact upon request.

### Experimental Model and subject details

#### Cell Culturing and differentiation

Two different batches of hiPSC-CMs were used for this study, a commercial one (Pluricyte® CMs provided by Ncardia, Gosselies, Belgium) and a laboratory handle one. The latter was produced by directed cardiac differentiation of previously generated control hiPSCs (Di Pasquale et al., 2013; Lodola et al., 2016, 2019b).

In brief, cardiac induction and generation of hiPSC-CMs were obtained as previously reported, using a chemically-defined serum-free protocol, which is based on activation (CHIR99021) and inhibition (IWR1) of the Wnt pathway in RPMI-B27 medium (Lian et al., 2013; Nakahama and Di Pasquale, 2016; Salvarani et al., 2019). In this work, CMs were differentiated from two different control iPSC lines (one male and one female) between the 18th and the 35^th^ passage in culture. Importantly, employed cell lines were regularly tested for being free of major chromosomal abnormalities by karyotype analysis. CMs were used for the experiments 25–30 days after spontaneous contracting activity was started, a differentiation stage at which they expressed the repertoire of sarcomeric proteins, calcium regulators, and ionic channels necessary for their correct functionality. Purity of differentiated CM populations was regularly checked before each experiment to be greater than 90% (not shown). The commercial batch was thawed and cultured following the manufacturer’s protocol. In terms of plating density, cells for both batches were seeded onto the material at a confluence comprised between 50% and 60% on fibronectin coated (15 μg/ml in PBS buffer solution) 18-mm round glass coverslip (VWR, Radnor, USA). The cells were maintained in incubator at 37°C and 5% CO2.

### Method Details

#### Ziapin2 Synthesis and uptake process

Ziapin2 was synthetized as previously described (DiFrancesco et al., 2020; Paternò et al., 2020b; Vurro et al., 2021). Micromolar concentrations (5 or 25 μM) of the compound were added to hiPSC-CMs cultures directly into the culture medium. Subsequently the cells were placed in the incubator at 37°C and 5% of CO2. After 7 minutes the medium was gently washed out and the petri dish was rinsed with fresh extracellular solution.

#### Electrophysiology

Standard whole-cell patch clamp recordings were performed on isolated cells with an Axopatch 200B (Axon Instruments) coupled with a Nikon Eclipse Ti inverted microscope. Acquisitions were performed with freshly pulled glass pipettes (4-7 MΩ), filled with the following intracellular solution [mM]: 12 KCl, 125 K-Gluconate, 1 MgCl2, 0.1 CaCl2, 10 EGTA, 10 HEPES, 10 ATP-Na2. The extracellular solution contained [mM]: 135 NaCl, 5.4 KCl, 5 HEPES, 10 Glucose, 1.8 CaCl2, 1 MgCl2. Acquisitions were performed with pClamp-10 software (Axon Instruments). Membrane currents were low pass filtered at 2 kHz and digitized with a sampling rate of 10 kHz (Digidata 1440 A, Molecular Devices).

A cyan excitation light (λ_ex_ = 470 nm) was provided by a LED system (Lumencor Spectra X) coupled to the fluorescence port of the microscope through a liquid-fibre. The photoexcitation density was 80 mW/mm^2^, as measured at the output of the 40X microscope objective (Pobj). Hyperpolarization and depolarization were measured as the minimum and maximum voltage, respectively, reached within 350 ms from the light-onset. Data were analyzed with Origin 9.0 (OriginLab Corporation) and MATLAB (MathWorks).

#### Capacitance recordings

Capacitance measurements were performed as previously described (Vurro et al., 2020, 2021). Briefly, a double sinusoidal voltage clamp signal was applied to the cell in whole-cell configuration. The response current signal was acquired, and membrane capacitance and resistance were extracted fitting the current with a custom MATLAB program. The capacitance value was extracted in dark condition and during light stimulation (20 ms pulse). The shorter pulse was used in order to consider mainly the effect related to Ziapin2 photoisomerization.

#### Intracellular Ca^2+^ measurements

Intracellular Ca^2+^ measurements were performed with a Nikon Eclipse TE200 microscope, and the extracellular solution was the one used for the electrophysiology measurements. Ca^2+^ dynamics were recorded in hiPSC-CMs incubated with 2 μM Fura-2 AM probe (Ex: 340/380 nm; Em: 510 nm) for 30 minutes and washed out for 5 minutes with extracellular solution. Ziapin2 (474 nm at power density of 273 μW/mm^2^) and Fura-2 AM excitations (340 and 380 nm, respectively at power density 63 and 173 μW/mm^2^) were provided using the same arc lamp light source coupled with high-speed OptoScan monochromator that allowed us to accurately select the wavelength. To collect local signals only (kinetics not distorted by propagation), a diaphragm in the optical path was adjusted to delimit a region of interest (ROI) enclosing single cells. The intracellular concentration of Ca^2+^ ([Ca^2+^]i) was monitored by measuring, for each ROI, the ratio of the mean fluorescence emitted at 510 nm when exciting alternatively at 340 and 380 nm (shortly termed ‘‘ratio’’), four to five ROIs were analyzed in each plate. An increase in [Ca^2+^]i causes an increase in the ratio. The following parameters were extracted from each CaT: diastolic Ca^2+^ (CaD), CaT amplitude (Amp, calculated as CaT peak—CaD), CaT rise-time (t_peak_) and CaT decay kinetics (τ_decay_).

#### Acquisition and analysis of hiPSC-CMs contractile behaviour

The Nikon Eclipse Ti inverted microscope described before was used to stimulate (Lumencor Spectra X, λ_ex_ = 470 nm, 20X objective, Pobj = 30 mW/mm^2^) and acquire video of the CMs contraction a frequency. An objective with a lower magnification was used to increase the amount of recorder cells resulting in a more robust analysis. Consequently, the resulting power density was lower due to the bigger area.

Contractile behavior was analyzed using a custom-built algorithm, implemented in MATLAB R2020a. The approach is based on the contraction-induced retraction of cell body towards nucleus and is effective even if there is no displacement of cell extremities or if the cellular edge cannot be detected properly.

The user defines one or more regions of interest (ROI) using bounding boxes. Each bounding box should contain one single cell. For each ROI a set of features is identified and tracked across video frames by means of Kanade-Lucas-Tomasi algorithm (Lucas and Kanade; Jianbo Shi and Tomasi, 1994). Since the ROI is delimiting a cell, the tracked features are expected to belong to cell body. Averaging the estimated motion fields of all these feature points returns a mean geometric transformation, which can be applied to the bounding box delimiting the ROI. The area of this bounding box is measured over time and presents minima at cellular contractions. The number of minima per time interval yields an estimate of the cell contraction rate. Contraction amplitude is measured as percent reduction of bounding box area at contraction and averaged over all contractions within a specific running observation window (e.g., pre). The box area derivative when cells contract evaluates cell contraction velocity. Data were analyzed with Origin 9.0 (OriginLab Corporation).

### Statistical Analysis

Data were expressed as means ± sem. Normal distribution was assessed using D’Agostino-Pearson’s normality test. To compare two sample groups, either the Student’s t-test or the Mann-Whitney U-test was used. To compare more than two sample groups, one-way ANOVA or Kruskal-Wallis were used. p < 0.05 was considered statistically significant. The analysis was carried out using Origin (OriginLab) and Prism (GraphPad).

## Supplementary

**Figure S1.**
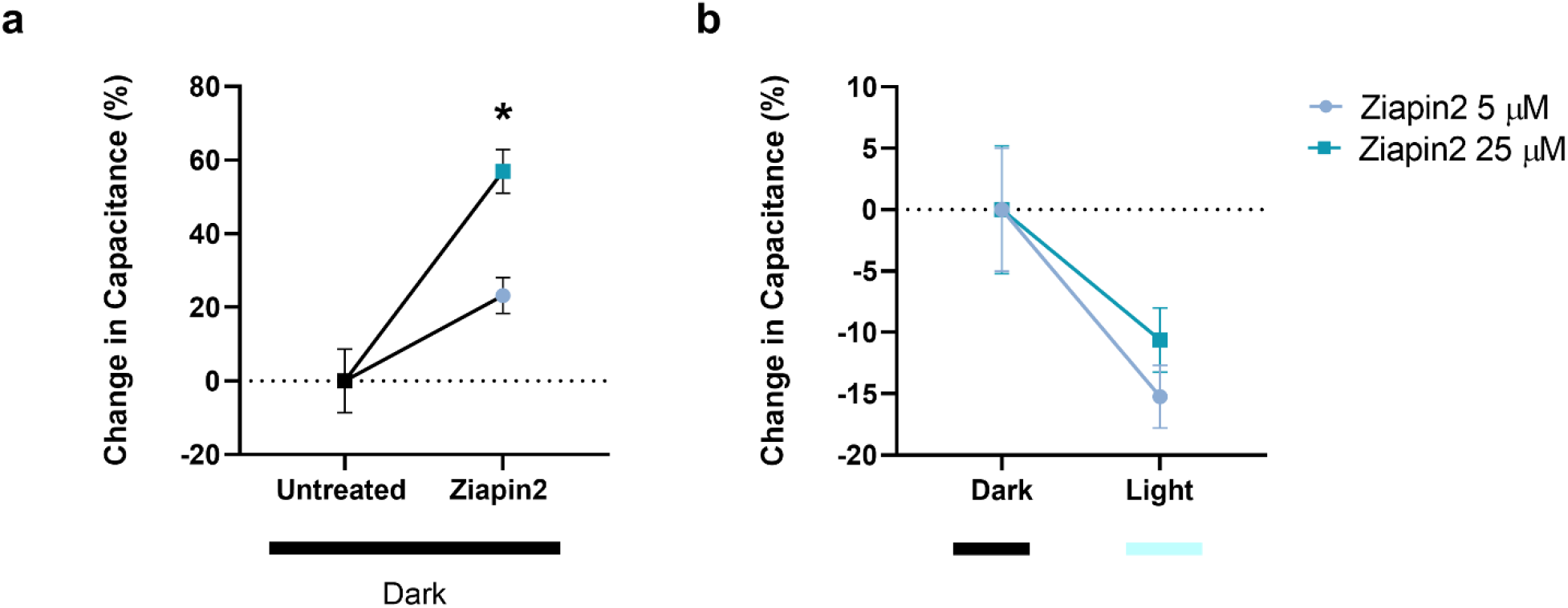
Ziapin2-mediated modulation of membrane capacitance. Evaluation of capacitance changes in hiPSC-CMs upon Ziapin2 membrane partitioning (**panel a**) and photoisomerization (**panel b**). Ziapin2 5μM represented with light blue dots while 25 μM in teal squares. Light power density, 80 mW/mm^2^. The experiments were carried out at room temperature (24°C). n = 15 and 20 for untreated and Ziapin2-loaded cells respectively from three independent differentiations. Data are represented as mean ± sem.

**Figure S2.**
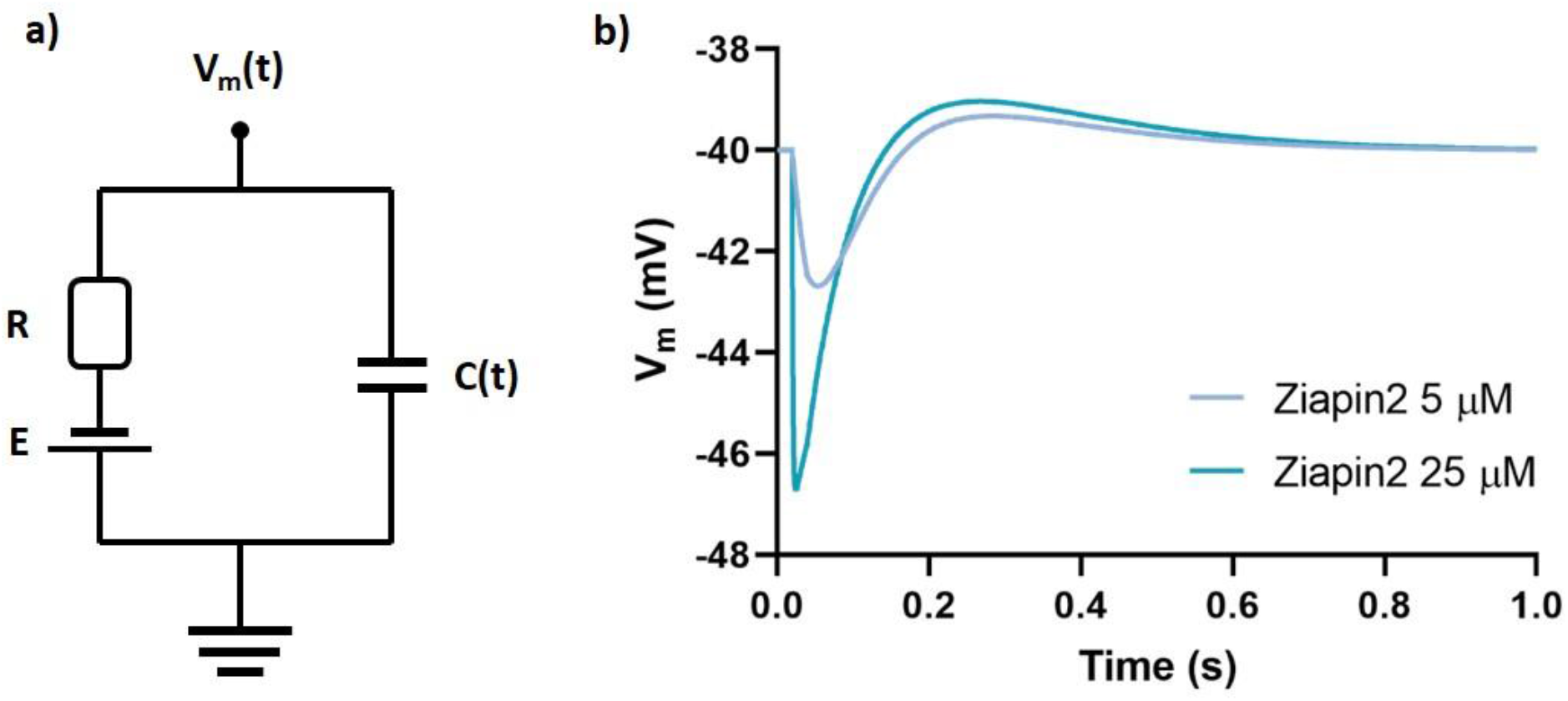
Simulation of membrane voltage. Simulated equivalent circuit (**panel a**) and capacitance-induced membrane potential variation (**panel b**). This simulation, performed as previously described (DiFrancesco et al., 2020), mimicks the effect of a 20 ms light pulse on Ziapin2 photoisomerization (a capacitance variation) and subsequent membrane potential modulation. To reproduce the observed effect, only the photoisomerization characteristic time has been reduced (thyp = 35ms for 5mM and thyp = 1ms for 25mM). This could be consistent with the localization of Ziapin2 in membrane domains more prone to favour isomerization.

**Figure S3.**
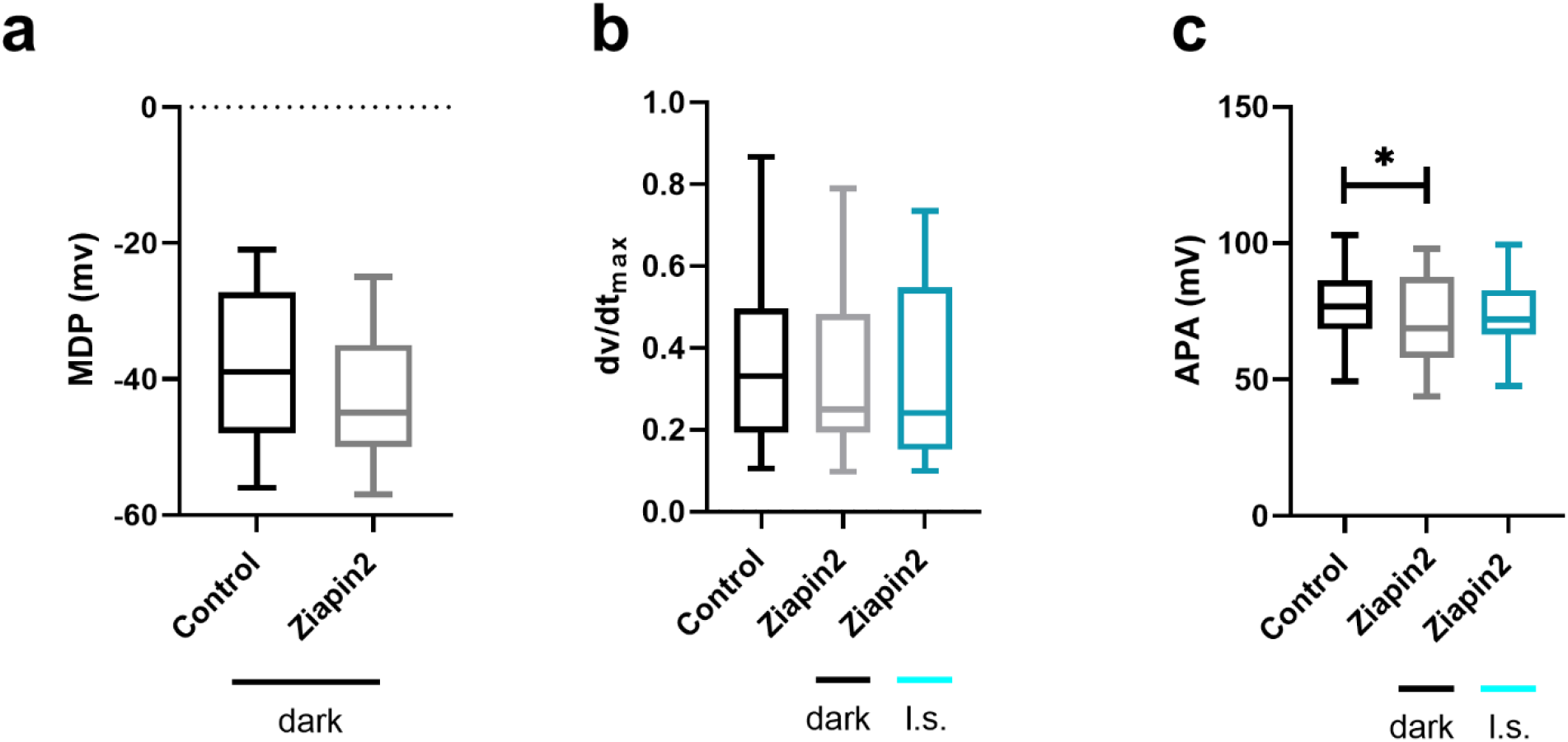
AP parameters in hiPSC-CMs exposed to Ziapin2. Comparison of the maximum diastolic potential (MDP, **panel a**), maximal rate of rise (dv/dtmax, **panel b**) and AP amplitude (APA, **panel c**) between spontaneous and light-evoked APs in 25μM Ziapin2 loaded hiPSC-CMs (n > 40 for each condition from three independent differentiations). Data are presented as box and whiskers plot (min to max). Light power density, 80 mW/mm^2^. The experiments were carried out at room temperature (24°C). * p < 0.05.

**Figure S4.**
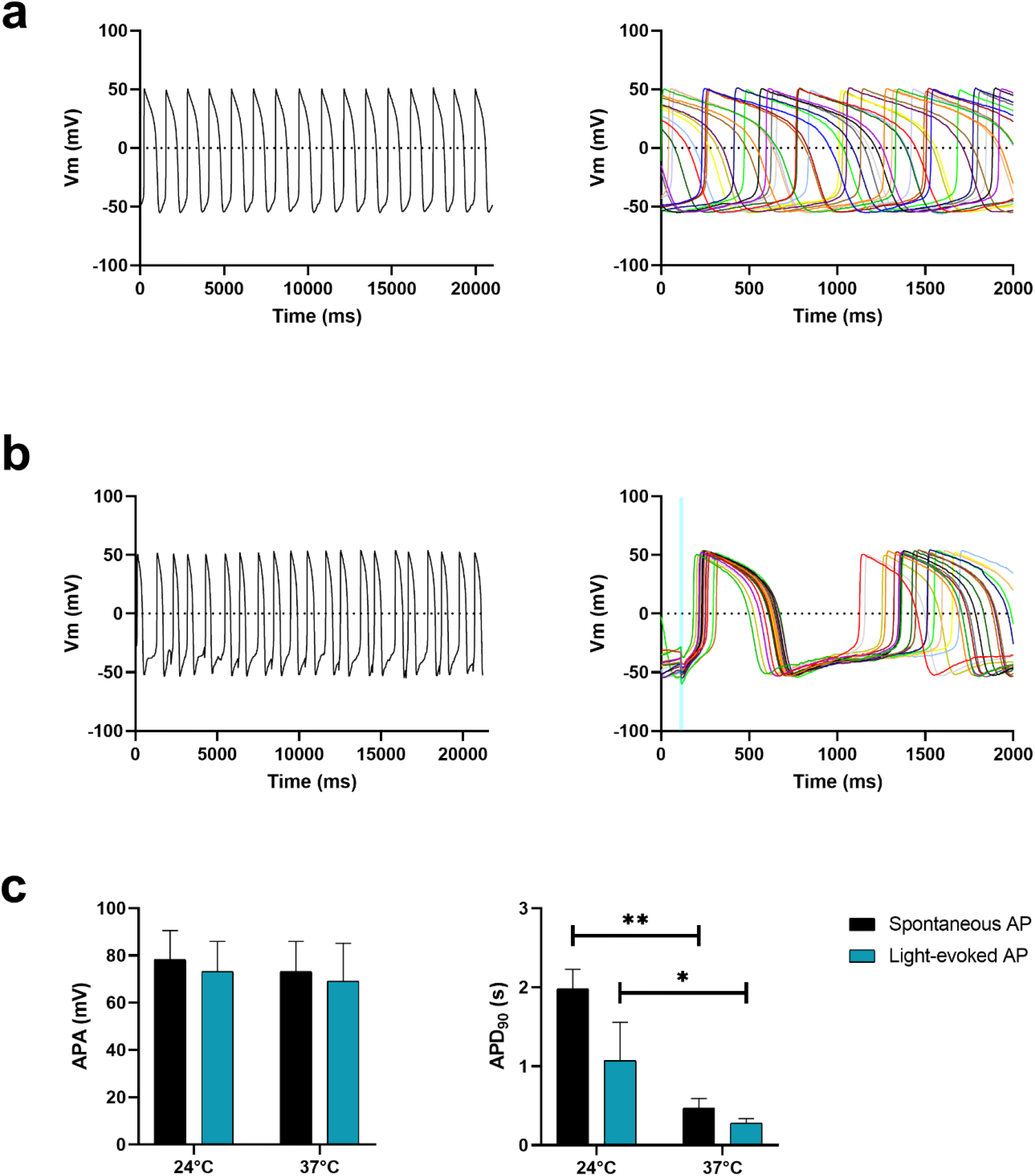
Light-induced V_m_ modulation in Ziapin2 loaded hiPSC-CMs at 37°C. **a)** Representative trace of spontaneous APs (left) and their distribution over a 2 second timescale (right). **b)** Representative trace of light-evoked APs (left) and their distribution over a 2 second timescale (right). The APs were generated in response to a light train stimulation (20 ms cycle length; 0.5 Hz stimulation frequency, represented as cyan shaded areas). Light power density, 80 mW/mm^2^. The experiments were carried out at 37°C. **c)** Comparison between spontaneous and light-evoked AP parameters (i.e., AP amplitude, APA and AP duration at 90% of repolarization, APD90) recorded either at room temperature or at 37°C (5<n<30, data are shown as mean ± sem). * p < 0.05, **p < 0.01.

**Figure S5.**
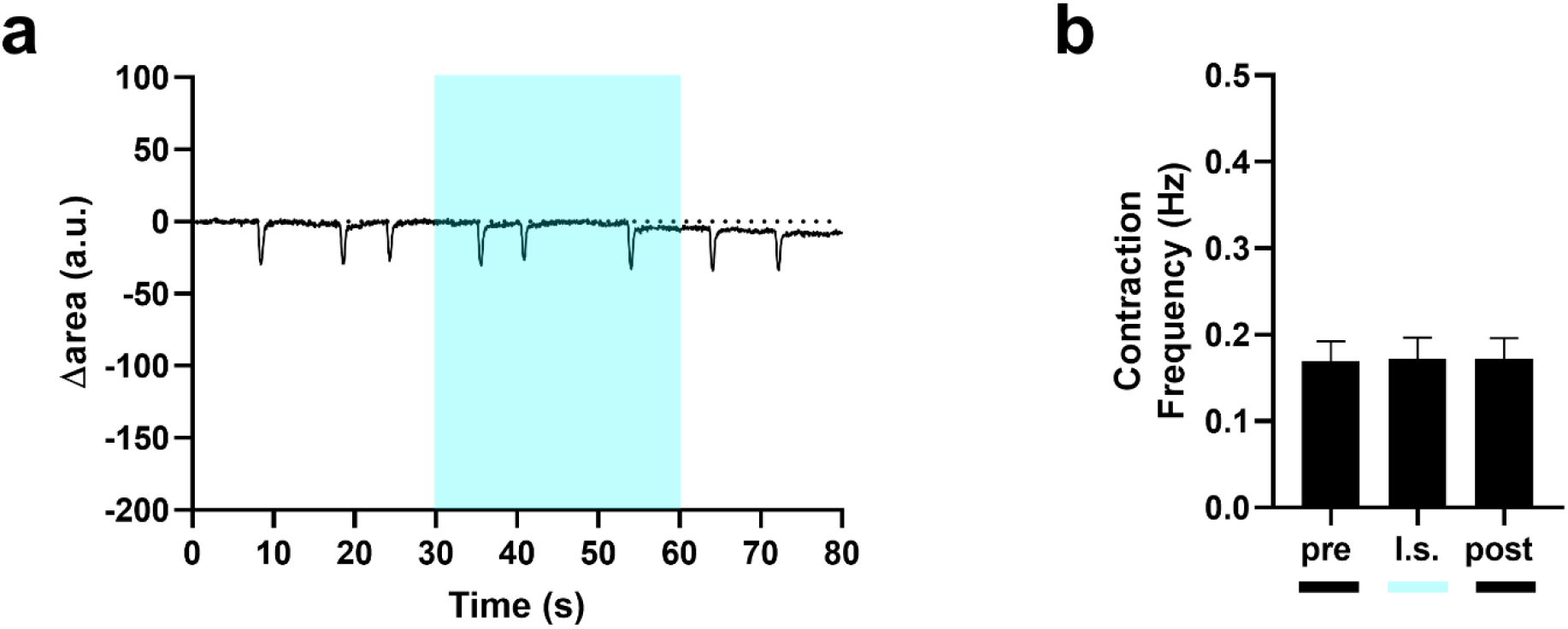
Light-induced contraction rate modulation in hiPSC-CMs incubated with vehicle. **a)** Representative trace of the contraction behaviour of DMSO-treated hiPSC-CMs. **b)** Contraction frequency before (pre), during (l.s.), and after (post) 1 Hz pulsed light stimulation at light power density = 30 mW/mm^2^ (n = 25 from three independent differentiations, data are shown as mean ± sem). The experiments were carried out at room temperature (24°C).

